# Exploring the role of mass immunisation in influenza pandemic preparedness: a modelling study for the UK context

**DOI:** 10.1101/775270

**Authors:** Luca Grieco, Jasmina Panovska-Griffiths, Edwin van Leeuwen, Peter Grove, Martin Utley

## Abstract

Existing modelling work on preparedness to pandemic influenza has focused on evaluating specific countermeasures for pandemics with specific characteristics (typically based on historical instances). The aim of this study was to inform policy on preparedness planning for pandemic influenza based on the assessment of a wide range of scenarios and free from restrictive assumptions about timing and features of the next pandemic.

We carried out epidemiological modelling and health economic analysis of an extensive set of scenarios, each comprising a combination of pandemic, vaccine and immunisation programme characteristics in presence or absence of access to effective antivirals. Preparedness policies that incorporate mass immunisation were evaluated on the basis of there being a given chance of a pandemic each year. To support understanding and exploration of model output, an interactive visualisation tool was devised and made available online.

We evaluated over 29 million combinations of pandemic and policy characteristics. Preparedness plans incorporating mass immunisation show positive net present value for a wide range of pandemic scenarios, predominantly in the absence of effective antivirals. Plans based on the responsive purchase of vaccine have greater benefit than plans reliant on the purchase and maintenance of a stockpile if immunisation can start without extensive delays. This finding is not dependent on responsively purchased vaccine being more effective than stockpiled vaccine, but rather is driven by avoiding the costs of storing and replenishing a stockpile.

While emerging technologies for rapid vaccine development and production increase the prospects for mass immunisation to be an effective countermeasure, policies based on the responsive purchase of vaccine not tailored to the pandemic should be explored. Focus is also required on the pandemic intelligence, decision, contractual and logistical processes on which timely commencement of immunisation in a pandemic is reliant.

## Introduction

The occurrence and impact of four major influenza pandemics in the last century (1918, 1957, 1968 and 2009 [1–3]) illustrate the threat to societies from pandemic influenza and the need for thorough preparedness planning to enable an effective response. Preparedness measures can include plans for immunisation programmes, plans for administering antiviral drugs that can reduce the duration and severity of infection, and agreeing the nature and trigger points for social distancing measures such as full quarantine, school, workplace and medical facility closure or travel restrictions [4–6]. This paper describes research done on behalf of the UK government to inform preparedness planning for a future influenza pandemic, specifically with regard to the role of mass immunisation.

Determining whether an intervention should be part of preparedness plans for a future, uncertain threat to public health is different from assessing an intervention in the context of an immediate, known threat.

Immunisation is an effective and cost-effective countermeasure to many infectious diseases [7]. In the context of an influenza pandemic however, the benefits of immunisation strategies are less clear; vaccines induce a narrow and strain-specific immunity [8] and, because pandemics occur only when there is marked shift in the strain of influenza circulating among humans, one cannot plan to have large volumes of a vaccine tailored to the pandemic strain available at the start of a pandemic.

Currently, there are two main preparedness options for mass immunisation in high-income settings. One is to maintain a stockpile of influenza vaccine that can be deployed early in a pandemic but which is not tailored to the pandemic strain, and hence likely to be less effective. The other is to negotiate in advance with manufacturers an option to purchase large quantities of a vaccine tailored to the pandemic strain but available later in the pandemic [4].

Initiatives aimed at supporting innovation and capacity for vaccine development and production offer the prospect of shortening the delay between a pandemic becoming evident and the availability of a vaccine [9] but the trade-offs between the timeliness of a mass immunisation programme and the efficacy of the vaccine used are complex.

Mathematical and computational approaches that model the spread of infection among a population have been used in numerous studies to evaluate the costs and benefits of different combinations of counter measures against pandemic influenza. Baguelin *et al.*, Prosser *et al.*, Ferguson *et al.* and Lugnér *et al.* [10–15] assessed the economic outcomes of vaccination strategies against pandemic influenza. Previous models have also explored antiviral treatment and immunisation strategies in parallel (e.g. Lee *et al.* [15], Newall *et al.* [16] or Khazeni *et al.* [17,18]). Recently, Halder *et al.* [19] investigated the cost-effectiveness of responsive purchase vaccination, taking into account a 6-month delay in vaccine availability, with and without combined social distancing and antiviral interventions.

This existing modelling work has largely been focused on evaluating specific countermeasures in the context of a pandemic with specified characteristics (in terms of infection spread and severity), typically reflecting an historic instance. However, policy decisions about preparedness plans need to be made without knowledge of the characteristics or timing of the next pandemic. For this reason, good preparedness plans are those that provide sufficient benefits at acceptable costs across a wide range of plausible future scenarios. At the request of colleagues at the UK Department of Health and Social Care, we worked to identify the circumstances under which preparedness plans involving mass immunisation would be considered good policy options. Our intent was to inform policy makers on the role of mass immunisation within pandemic preparedness planning and the extent to which potential improvements in vaccine development and production could enhance policies based on the responsive purchase of vaccine.

## Materials and methods

We adapted an existing epidemiological model of influenza spread among the UK population to enable the evaluation of a large number of scenarios, each characterised by a unique combination of: the features of a mass immunisation programme, the nature of the next influenza pandemic and the availability or otherwise of effective antiviral drugs with which to treat infected cases. For each scenario, we used the output of the epidemiological model in a health economic analysis to estimate the net benefit of mass immunisation in that scenario. Given the very large number of scenarios explored, we devised a compact visualisation of the model output to enable insights to be drawn about different preparedness policies. We describe these components of our work below.

### Epidemiological model

We conducted a scoping review of the development and use of models to assess the cost-effectiveness and net-benefit of different mass immunisation strategies against pandemic influenza (see Supplementary File S1 for details). Of those models implemented and available in the public domain, we chose to adapt and use that of Baguelin and van Leeuwen [11,20]. This epidemiological model, which is based on a system of ordinary differential equations, was designed to estimate the number of influenza susceptible, exposed, infected and recovered individuals over time for the purpose of evaluating countermeasures to *seasonal* influenza. It was recently made publicly available [20] in the open source programming language R (https://www.r-project.org). In adapting it for our purpose we took out the seasonality, simplified the age structure used within the model, and wrote a “shell script” to implement and collate the output from a large number of “model runs”. Liaising closely with the authors of the model (one of whom joined the study team), we devised a way to estimate the impact of mass immunisation alone and in combination with the distribution of effective antivirals for treating infected cases. See Supplementary File S2 for details of our model adaptations.

The adapted model was set up with some fixed parameters: the size of the UK population (65,300,000), the average number of contacts per day per individual (13) and the number of infected individuals at the time point considered to be the “start of the pandemic” within the model (2,000). It took as input (Table 1) some variables reflecting the features of an immunisation programme and others reflecting the nature of the pandemic to be modelled. The model gave as output the estimated number of susceptible, exposed, infected and recovered individuals each day for a year following the start of a pandemic, as well as the estimated number of hospitalisations and the number of deaths associated with the infections.

**Table 1.** Ranges of parameter values.

| Type | Parameter | Range of parameter values used in analysis |
| --- | --- | --- |
| Features of the pandemic to be modelled | Basic reproduction number ($R_0$) | {1.2, 1.4, ..., 2.4} |
|  | Case fatality ratio (CFR) | {0.04%, 0.2%, 2%} |
|  | Susceptibility of the population | {80%, 90%} |
|  | Probability of second pandemic | {0, 0.05, 0.1} |
| Features of an immunisation programme | Vaccine efficacy | {20%, 40%, 60%, 80%} |
|  | Uptake of vaccination among the population | {5%, ..., 100%} |
|  | The time between the start of the pandemic and the start of immunisation programme | {0, 1, 2, ..., 22} weeks |
|  | Time over which vaccine coverage is achieved | {2, 4, 8, 12, 16} weeks |
|  | Vaccine shelf-life | {1, 2, 5} years |
|  | Affordability threshold | £ {0.5bn, 1bn, 1.5bn, 2bn} |

Parameters describing different features of the pandemic and immunisation programme modelled, together with the range of values used in the analysis.

Another characteristic of each scenario was the chance of there being a further set of influenza infections once the initial pandemic has petered out. Note that these further infections were not modelled as a “second wave” within a single run of the epidemiological model. Rather, we used the model to calculate the number of additional cases that would be associated with a second pandemic with a moderately high basic reproduction number (R_0_ = 2.2), a moderate case fatality ratio (CFR = 0.2%), lower population susceptibility (50%) and with any additional immunity due to mass immunisation during the first wave assumed to be in place at the start of this second pandemic (cf. Supplementary File S2).

### Health Economic analysis

For each combination of the programme and pandemic variables given in Table 1, the epidemiological model was run 4 times: once with no countermeasures, once with only the specified mass immunisation programme, once with only the counter-measure of distributing effective antivirals to infected individuals and once with both of these counter-measures. From these we derived the health benefits of the specified immunisation programme in the presence or absence of effective antivirals. We did this by calculating the Quality-Adjusted Life Years (QALYs) gained from avoided clinical influenza cases, hospitalisations and deaths attributable to immunisation, using a discount rate of 1.5% for health benefits over a time horizon of 10 years. We then converted these discounted health benefits to a monetary value based on the monetary value of a QALY used by the UK Government. For each scenario, costs were calculated for each of two preparedness policies for immunisation. Within the model, policies to stockpile vaccine incur purchase, wastage and storage costs every year and distribution and administration costs only in the event of a pandemic. Policies to purchase vaccine responsively incur purchase, distribution and administration costs in the event of a pandemic plus the cost of an annual fee payable every year to the manufacturers for the option to buy large quantities of vaccine. Under each policy, we calculated the total discounted cost of the policy, using a discount rate of 3.5% for monetary costs with a time horizon of 10 years, with the key assumption that the annual chance of a pandemic actually happening is 3% [21]. We then calculated the Net Present Value (NPV) for the policy, defined as the discounted monetised health benefits minus the discounted costs. We also assessed the upfront costs associated with a preparedness policy against a threshold value to reflect the possibility that some policies can be deemed unaffordable even if cost-effective. The details of the health economic analysis can be found in Supplementary File S2. We report the list of fixed parameter values used in Supplementary File S3.

### The scenarios explored and presentation of results

The value ranges and increments used for each programme and pandemic variable within the epidemiological model and health economic evaluation are shown in Table 1. Overall, we ran 29,211,840 combinations of programme and pandemic variables. For reference only, we note that estimated R_0_ for the 2009 and 1957 pandemics were in the ranges [1.3, 1.7] and [1.47, 2.27], respectively [22], and estimated CFR for the same pandemics were in the ranges [0.01%, 0.08%] and [0.74%, 1.8%], respectively [19].

As our purpose was not to evaluate the role of antivirals, we took a simplistic approach to characterising the use of antivirals in our evaluation of immunisation. When effective antivirals were assumed to be deployed the modelled impact (a reduction in infection duration and a reduction in case fatality ratio for infected cases) was the same whatever the characteristics of the pandemic strain.

We constructed “heat-map” graphs to display the net present value (NPV) of a particular preparedness policy (pre-purchase of vaccine or responsive purchase of vaccine) in the presence or absence of effective antivirals (Figure 1). In these graphs, colour-coded regions identify the combinations of variables under which a preparedness policy has negative net present value (shaded pink) or positive net present value (darkening shades of green with increasing NPV). Greyed out regions of these graphs identify combinations of variables for which the cost of the policy exceeds the given affordability threshold. We arranged the graphs as arrays of “tiles” with each tile showing how NPV varies with the coverage of the immunisation programme and the number of days between the start of the pandemic and the start of the immunisation for a specific set of other variable values, some of which change from tile to tile within the array.

**Figure 1.**
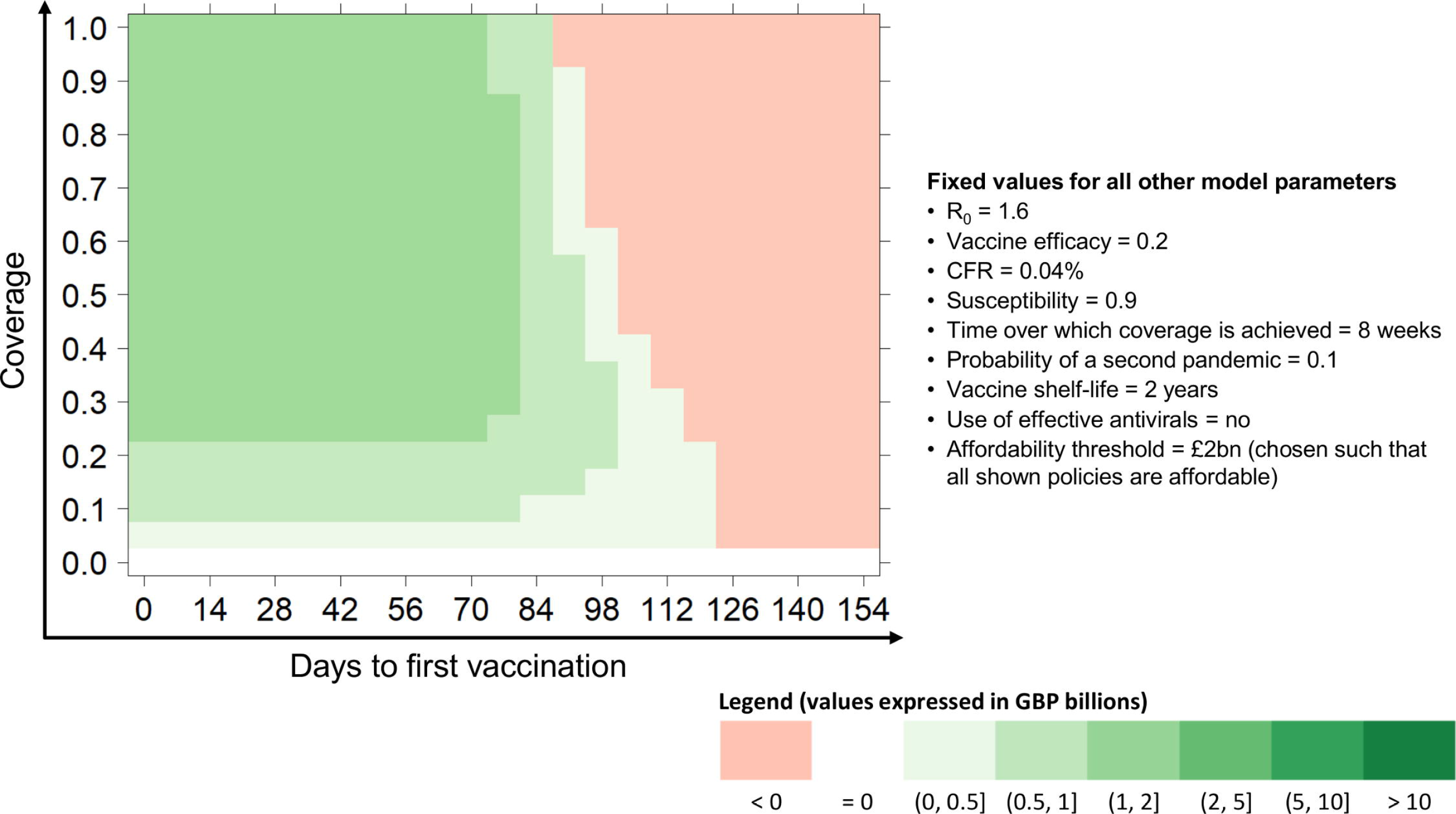
Heat-map graph displaying policy’s net present values. We show the NPV of a given policy (in this example: pre-purchase strategy, no antivirals) as a function of days to first vaccination (time lag between the start of the pandemic and the start of the immunisation programme) and coverage (uptake of vaccination among the population), with all remaining model parameters fixed. Colour-coded regions identify the combinations of variables under which the policy has negative net present value (shaded pink) or positive net present value (darkening shades of green with increasing NPV).

To allow visualisation and exploration of the full set of model runs and analyses constructed, we embedded our programming code into an interactive webtool using a service provided by shinyapps.io (https://www.shinyapps.io/). The tool (accessible at https://vaccinparamspaceanalysis.shinyapps.io/shinyPlots/) enables users to reproduce our analyses and compare up to 4 heat-map graphs at a time. All possible combinations of parameters listed in Table 1 can be explored by selecting them in the sliders and dropdown lists provided.

### Sensitivity analysis

We tested the sensitivity of our results to variations in the parameter values used in the Health Economic analysis. For each of a subset of the heat-map graphs shown in the Results section, we varied in turn each of the Health Economic parameters by +/−10% of their original value, recalculating the corresponding NPVs and constructing new heat-map graphs for visual comparison with the original graph.

## Results

The full set of results can be explored using the tool (https://vaccinparamspaceanalysis.shinyapps.io/shinyPlots/). As would be expected, the model output shows that, other things being equal, the benefits of immunisation increase with increasing efficacy of the vaccine and increasing case-fatality of the pandemic. Note that we do not report here absolute values of net present value as these vary from scenario to scenario, but rather set out the range of scenarios where different policies have positive net present value.

We summarise below some key findings of interest for those faced with decisions about whether to include plans for mass immunisation as part of national pandemic preparedness policy. In terms of affordability, policies based on use of a pre-purchased vaccine incur upfront costs estimated to exceed £1bn if target coverage exceeds around 60% of the UK population.

### Value of mass immunisation as a lone counter-measure

If a nation does not have a stockpile of antivirals, or considers that antivirals will not be effective against the range of pandemics they want to prepare for, or considers timely distribution of antivirals to infected cases in a pandemic infeasible, preparedness plans for mass immunisation have positive net present value in a wide range of circumstances.

A policy for mass immunisation using a pre-purchased vaccine with low efficacy (20%) has positive net present value unless any pandemic occurring is mild in terms of case fatality and slow spreading (Figure 2 (a)) or more rapidly spreading with a substantial delay before starting the immunisation programme (Figure 2 (c-e)(i-j)). As an aside it is worth noting that, for such policies, if any pandemic occurring is mild there is a point at which increasing target coverage reduces net benefit, in certain cases to the extent where there is a net-loss (Figure 2 (a-e)).

**Figure 2.**
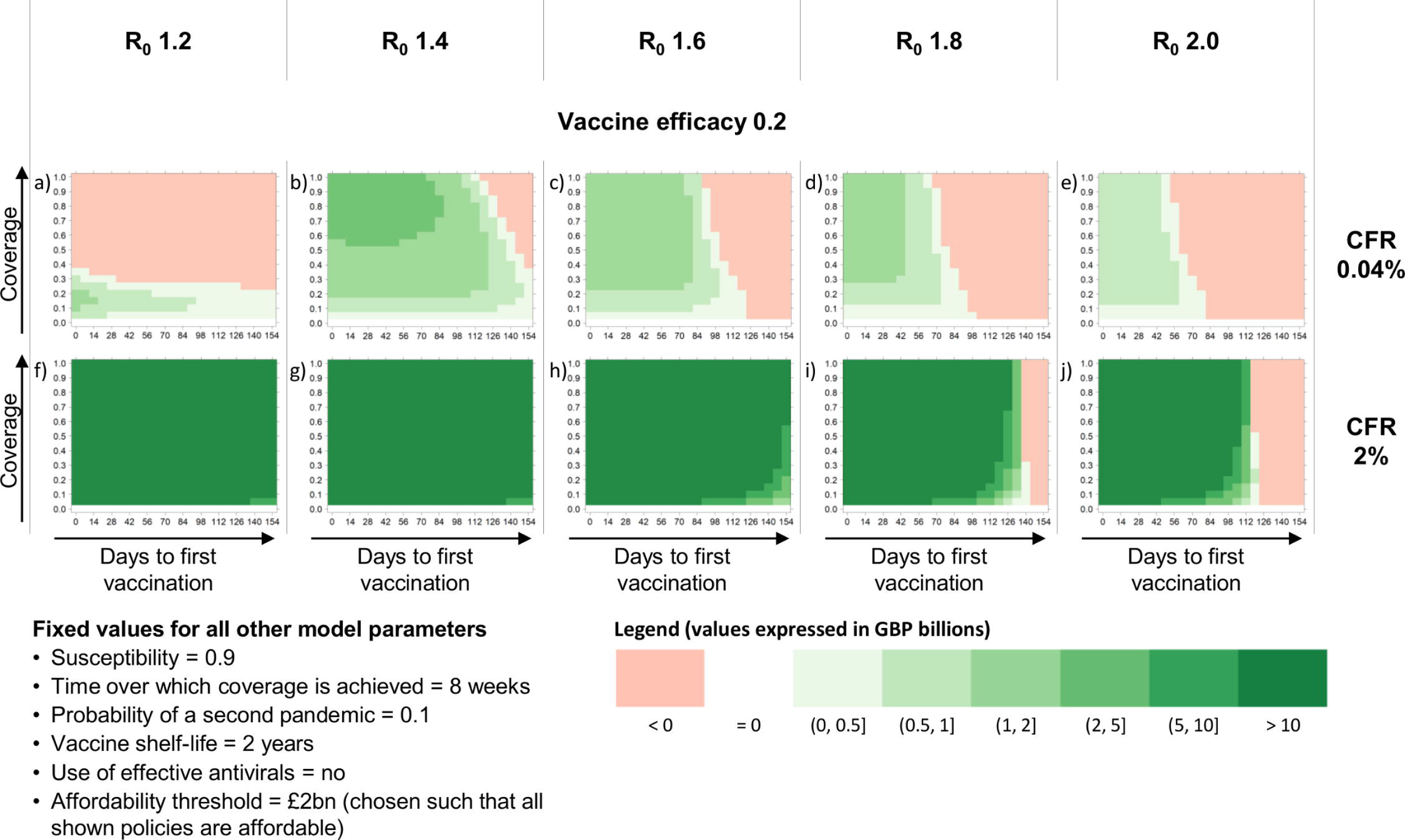
Pre-purchase strategy, no antivirals. Each heat-map shows, for a given combination of R_0_, vaccine efficacy and case fatality ratio, the net present value of this policy as a function of days to first vaccination (time lag between the start of the pandemic and the start of the immunisation programme) and coverage (uptake of vaccination among the population).

Figure 3 shows that a policy for mass immunisation with responsive-purchase vaccine has positive net-benefit even with low efficacy (a-d)(g-j), unless the pandemic occurring is rapidly spreading and there is a delay in starting the immunisation programme of 3-4 months (e-f)(k-l).

**Figure 3.**
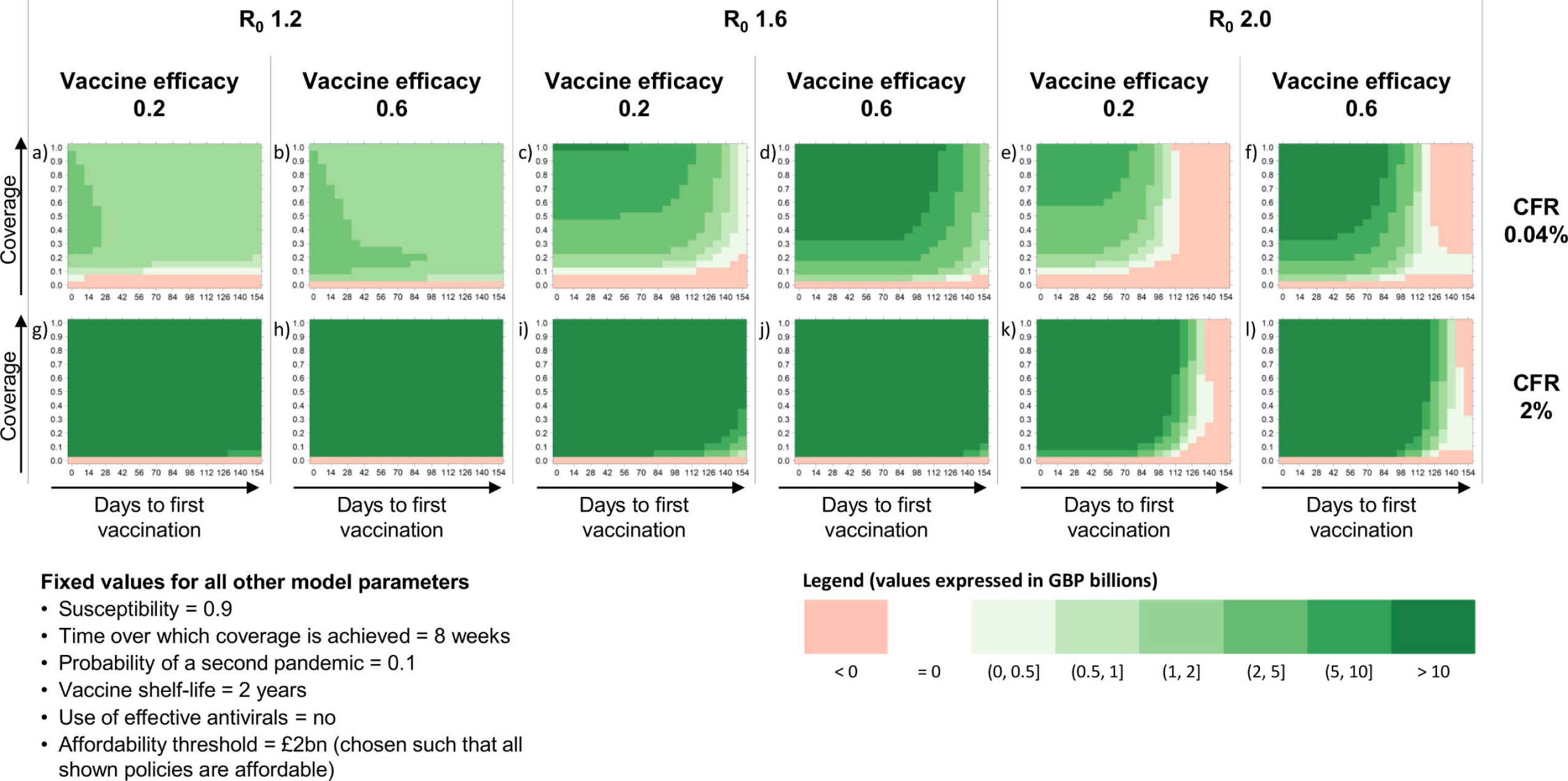
Responsive purchase strategy, no antivirals. Each heat-map shows, for a given combination of R_0_, vaccine efficacy and case fatality ratio, the net present value of this policy as a function of days to first vaccination (time lag between the start of the pandemic and the start of the immunisation programme) and coverage (uptake of vaccination among the population).

### Incremental value of mass immunisation to an effective policy of distributing antivirals

If a nation has a stockpile of antivirals, consider that they will be effective against the range of pandemics they want to plan for and that they can be deployed in a timely manner, our findings suggest that there are limited circumstances where an additional immunisation programme has positive net present value. If any pandemic occurring has low or moderate speed of spread (R_0_ ≤ 1.6), mass immunisation has negative net present value in parallel with antiviral use (Figure 4 (a)(e) and Figure 5 (a-b)(i-j)). If any pandemic occurring has R_0_≥1.8, pre-purchase of a vaccine with 20% efficacy only has positive NPV in parallel with effective antivirals if the pandemic has a high case fatality ratio (Figure 4 (b-d)(f-h)). A responsive-purchase policy can have positive NPV if a pandemic occurring has a lower fatality ratio but only if it would spread very rapidly in the absence of counter-measures (R_0_≥2) (Figure 5 (e-h)).

**Figure 4.**
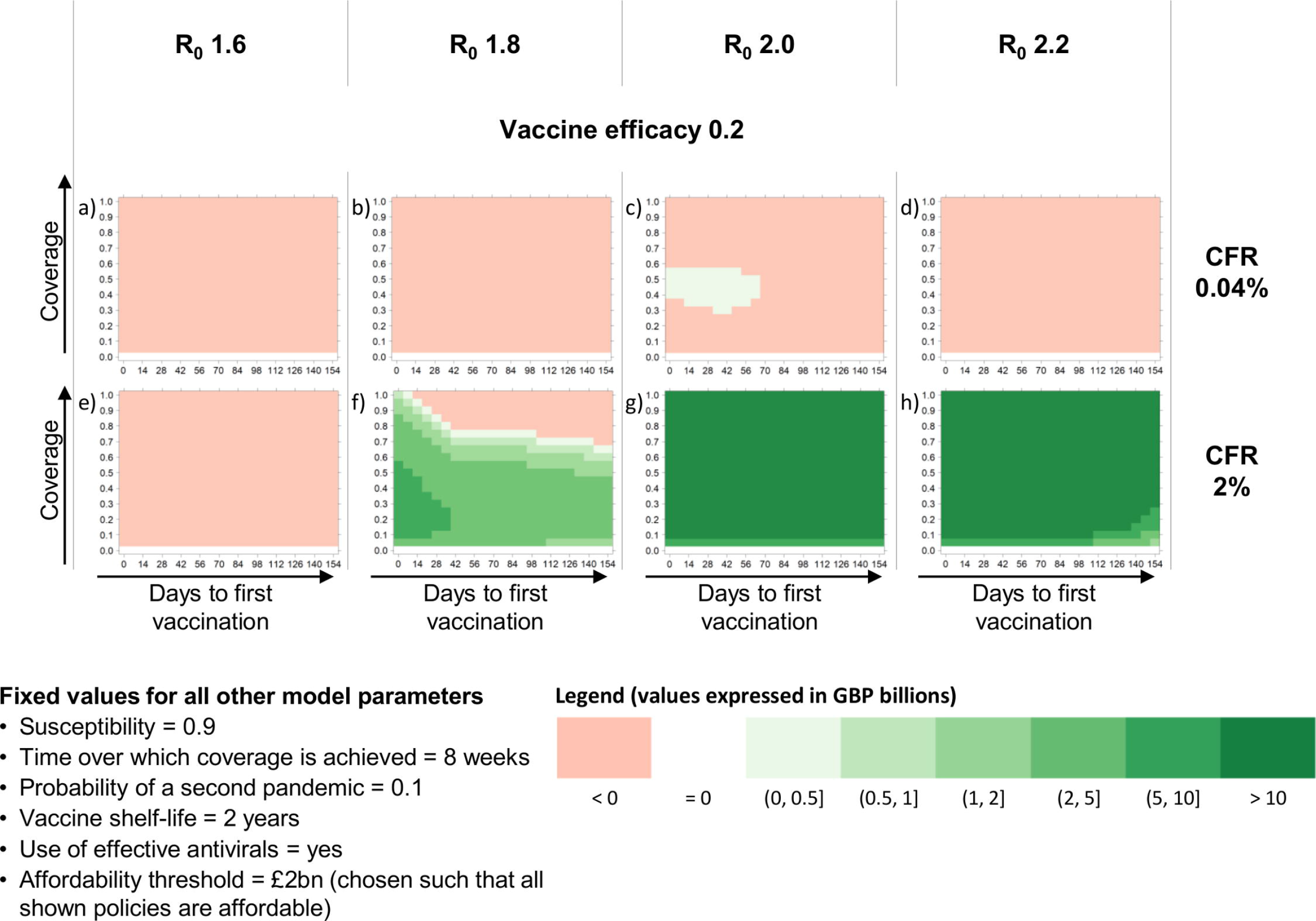
Pre-purchase strategy, with antivirals. Each heat-map shows, for a given combination of R_0_, vaccine efficacy and case fatality ratio, the net present value of this policy as a function of days to first vaccination (time lag between the start of the pandemic and the start of the immunisation programme) and coverage (uptake of vaccination among the population).

**Figure 5.**
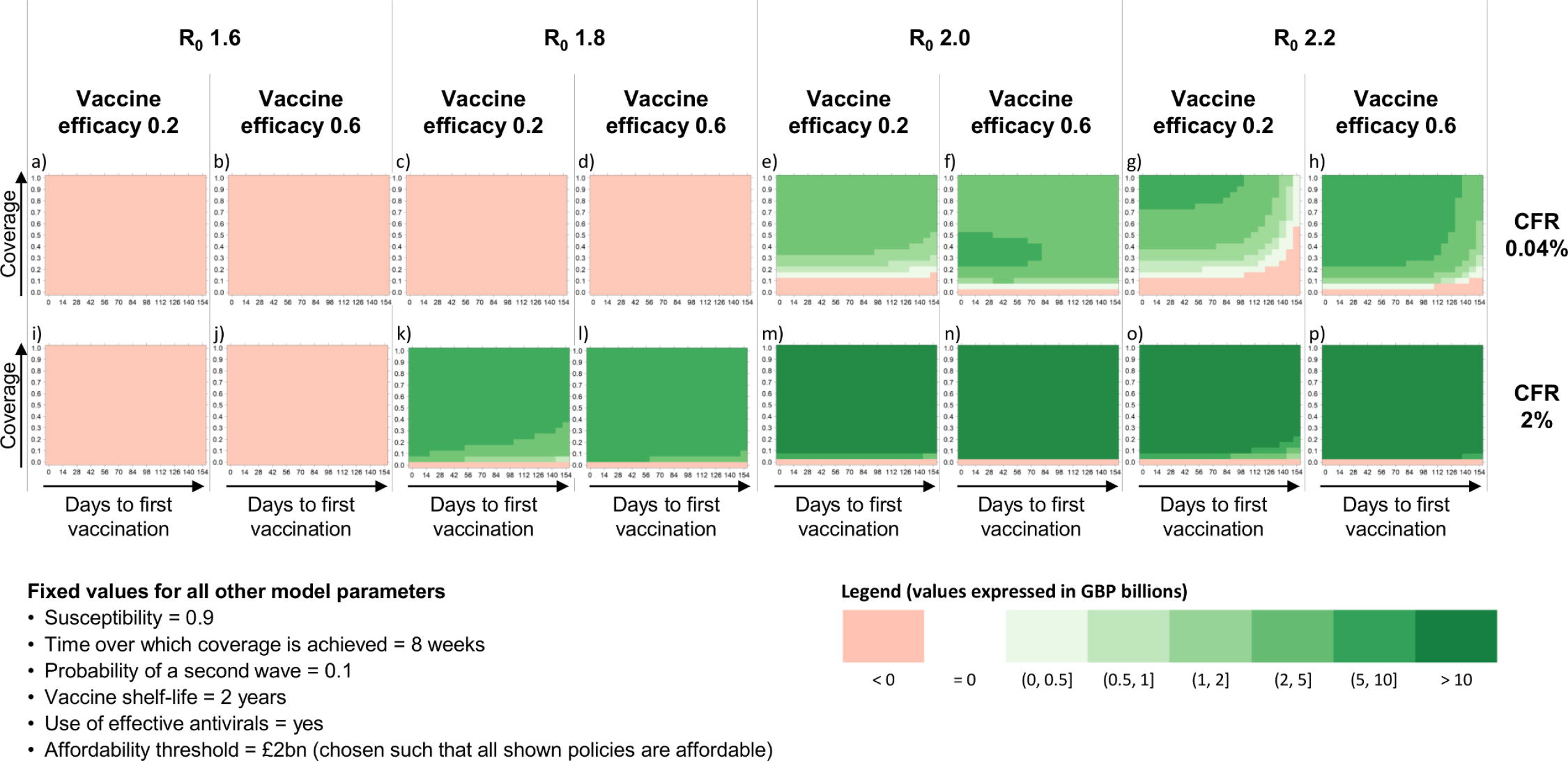
Responsive purchase strategy, with antivirals. Each heat-map shows, for a given combination of R_0_, vaccine efficacy and case fatality ratio, the net present value of this policy as a function of days to first vaccination (time lag between the start of the pandemic and the start of the immunisation programme) and coverage (uptake of vaccination among the population).

### Vaccine efficacy versus early programme start

In deciding between the preparedness options of stockpiling a pre-pandemic vaccine or paying to have the option to purchase a vaccine tailored to the pandemic strain, a comparison of tiles Figure 2(e) and Figure 3(f) and of tiles Figure 2(j) and Figure 3(l) is instructive. If the pandemic occurring is mild but spreads rapidly (CFR=0.04%, R_0_=2.0) pre-purchase of a vaccine with 20% efficacy offers modest net benefit if the programme can be started within 2 months of the pandemic while the potential advantages of deploying a more effective vaccine bought responsively are only realised if a programme can be started within 4 months (for 60% efficacy). For a higher case-fatality, any advantage of having a more effective vaccine through responsive purchase is dependent on immunisation starting within 5 months. This time sensitivity lessens for a pandemic with a moderate or low R_0_ (e.g. Figure 3(g-j)) or if immunisation happens in parallel with effective antivirals (e.g. Figure 5 (m-p)).

It is worth noting that responsive purchase can have advantages over stockpiling even if the vaccine bought responsively is not more efficacious (for instance Figure 3(c) compared to Figure 2(c) or 3(e) to 2(e)) if an immunisation programme based on responsive purchase can be started sufficiently quickly (4-5 months depending on speed of spread of pandemic). This is because responsive purchase avoids the cost of storing and replenishing a stockpile in the period up to the next pandemic.

### Sensitivity to Health Economic assumptions

In Supplementary File S4, we present the results of our sensitivity analysis carried out with respect to six of the tiles shown in Figures 2 to 5. For some of the parameters (e.g. QALY loss per death, pandemic probability, monetised QALY value) we can observe variation in the steepness of the contour lines represented in the tiles. However, the main findings reported in the previous sections are not sensitive to the 10% changes in health economic parameter values.

## Discussion

We have evaluated policies for influenza pandemic preparedness involving mass immunisation across a wide range of scenarios. This approach enables policy makers to assess pandemic preparedness policies of mass immunisation without having to predict the precise characteristics of the next pandemic.

Our results suggest that if a nation has a stockpile of antivirals that it is confident can be effectively deployed to treat infected cases in the advent of a pandemic, there are limited circumstances where a policy of an additional programme of mass immunisation has net benefit.

In the absence of effective antivirals or other countermeasures, a preparedness policy of mass immunisation has positive net benefit in a large range of circumstances. Overall, a strategy based on responsive purchase of vaccine in the event of a pandemic is beneficial in a wider set of pandemic scenarios than a strategy based on maintaining a stockpile of vaccine so long as the immunisation programme can be started sufficiently quickly. This finding was anticipated, but our model output allows policy makers to understand what counts as “sufficiently quickly” for the range of pandemic scenarios they decide to plan for. It is worth noting that, where responsive policies perform better than stockpiling, much of the advantages stem from avoiding the cost of storing and replenishing a stockpile, and are not heavily dependent on the vaccine deployed being more efficacious.

The epidemiological model and the health economic model used, the fixed parameters and the range of scenarios explored were all chosen in consultation with colleagues at the Health Protection Analytical Team at the UK Government’s Department of Health and Social Care to be aligned with current UK planning assumptions. This ensured that the model outputs would be relevant to decision processes in the UK, enhancing the utility of our research to our sponsor but, arguably, limiting its ready application to the policy contexts of other nations.

While a great many combinations of the model variables can be explored using the visualisation tool we constructed, some key assumptions within the model remain fixed and, as with any modelling study, there are limitations related to the validity of these assumptions. For instance, the annual chance of there being an influenza pandemic is taken as 3%. While in line with planning assumptions in the UK, this is an educated guess at best. Note that the higher the likelihood of a pandemic, the greater the anticipated benefit associated with preparedness policies incorporating mass immunisation. Another assumption that could be challenged is that the cost per dose of a vaccine to be stockpiled is assumed within our analysis to be the same as the cost per dose of a vaccine bought responsively. Also, we have not accounted for intrinsic limits on vaccine utilisation due to inevitable supply chain losses. Changes to these and other parameters and assumptions could be explored within the same analytical framework.

Decisions about policies for pandemic preparedness have to be made in the absence of knowledge about the timing and characteristics of the next influenza pandemic. Where other studies in this area [10–16,17,19] have focussed on evaluating countermeasures in the context of one or two specific sets of pandemic characteristics, the main strength of our approach is that, by exploring a vast range of different pandemic scenarios and by accounting for costs incurred in years when there is no pandemic, we can identify the range of circumstances under which a particular policy has net benefit and under which it does not. We consider that our approach is more attuned to the decisions that face policy makers and represents a direction of research that is necessary for better control of future influenza pandemics with currently unknown characteristics.

A limitation of our work compared to that of others is that, in focussing on the net present value of mass immunisation as a lone countermeasure or as an addition to the distribution of antivirals to infected individuals, we have not sought to identify the most effective combination of countermeasures. For instance, work by Newall *et al.* [16] and Khazeni *et al.* [18] suggests that expanded vaccination (mass immunisation) combined with effective antivirals use is the most beneficial for specified R_0_ values. Our model output (not shown) is consistent with this and extends the finding to other pandemic characteristics. Halder *et al.* [19] simulated both pre-emptive (i.e. pre-purchase) and reactive (i.e. responsive-purchase) vaccination strategies, combined with a range of social distancing and antiviral measures. They found that if pre-pandemic vaccines developed are less than 30% effective, the policy of pre-purchase vaccination is less cost effective than the responsive purchase vaccination strategy. Our work identifies the circumstances for which this is the case. We note that our results on the relative affordability of the pre-purchased compared to the responsive-purchase vaccine differ to those in Halder *et al.* [19]. Because they include social distancing prior to mass immunisation, the policies they evaluated incur considerable productivity losses that are not a feature of the policies we have evaluated.

One simplification in our work is that the epidemiological model we have used is not age and risk group stratified. We made this choice as we wanted to assess mass immunisation against future pandemics for which any age effects are unknown rather than assess programmes targeted at specific age/risk population cohorts or where there are strong assumptions about age-dependent susceptibility or case-fatality. As a consequence, the contact pattern we used is simplistic and other deterministic [10, 11, 14] or stochastic, agent-based [13, 16, 19] models would give more realistic predictions for the spread of a pandemic within and between different population cohorts for a strain with known age and risk group-dependent characteristics.

The results and the code underpinning the analysis have been shared in full with the Health Protection Analytical team at DH who are using them to inform policy on pandemic preparedness and policy related to new technologies for vaccine development and production. As a range of technologies emerge for improvements in vaccine development and production, our work could prove very useful in determining the trade-offs between timeliness and efficacy associated with different innovations in techniques for vaccine development and mass-production. Also, our findings point towards there being value in exploring polices that involve the responsive purchase of vaccine that is not tailored to the pandemic.

However, one question not addressed in this work relates to the speed with which a mass immunisation programme could or would be instigated in the event of a pandemic. For responsive purchase strategies, the time to develop and produce sufficient vaccine is currently the rate limiting step. It currently takes five to six months for an approved vaccine to become available after a new influenza virus strain is isolated [23, 24]. If evaluating scenarios where production is speeded up significantly, the times taken for the other decision making, contractual, and logistical processes involved in instigating an immunisation programme may need to be considered.

By design, the analysis conducted and the presentation of results do not account for the fact that some combinations of pandemic characteristics are more likely than others. Future work could incorporate elicitation of expert opinion to restrict the analysis to a smaller set of scenarios considered sufficiently plausible to plan for. Also, the framework could be adapted to explore more nuanced scenarios, for instance scenarios where there is considered to be a small chance of each of several strains with different specified characteristics.

## Acknowledgements

We would like to thank Prof Richard Pebody (Public Health England) and Dr Marc Baguelin (Public Health England and London School of Hygiene and Tropical Medicine) for helpful discussions related to this work.

## Supporting information

**Supplementary File S1.** Literature search of existing models that evaluate the cost-effectiveness of different vaccination strategies against pandemic influenza.

**Supplementary File S2.** Panovska-Griffiths J, Grieco L, van Leeuwen E, Grove P, Utley M. A method for evaluating the net-benefit of different preparedness planning policies against pandemic influenza. Methods X, under review.

**Supplementary File S3.** Economic model’s fixed parameter values used in our analyses.

**Supplementary File S4.** Sensitivity analysis results.

